# Taxonomic hypotheses and the biogeography of speciation in the Tiger Whiptail complex (*Aspidoscelis tigris*: Squamata, Teiidae)

**DOI:** 10.1101/2020.10.05.327270

**Authors:** Tyler K. Chafin, Marlis R. Douglas, Whitney J.B. Anthonysamy, Brian K. Sullivan, James M. Walker, James E. Cordes, Michael E. Douglas

## Abstract

(225)Biodiversity in southwestern North America has a complex biogeographic history involving tectonism interspersed with climatic fluctuations. This yields a contemporary pattern replete with historic idiosyncrasies often difficult to interpret when viewed from through the lens of modern ecology. The *Aspidoscelis tigris* (Tiger Whiptail) complex (Squamata: Teiidae) is one such group in which potential taxonomic boundaries have been confounded by a series of complex biogeographic processes that have defined the evolution of the clade. To clarify this situation, we first generated multiple taxonomic hypotheses, which were subsequently tested using mitochondrial DNA sequences (ATPase 8 and 6) evaluated across 239 individuals representing five continental members of this complex. To do so, we evaluated the manner by which our models parsed phylogenetic and biogeographic patterns. We found considerable variation among species ‘hypotheses’, which we suggest in part reflects inflated levels of inter-population genetic divergence caused by historical demographic expansion and contraction cycles. Inter-specific boundaries with *A. marmoratus* juxtaposed topographically with the Cochise Filter Barrier that separates Sonoran and Chihuahuan deserts (interpreted herein as case of ‘soft’ allopatry). Patterns of genetic divergence were consistent across the Cochise Filter Barrier, regardless of sample proximity. Surprisingly, this also held true for intraspecific comparisons that spanned the Colorado River. These in turn suggest geomorphic processes as a driver of speciation in the *tigris* complex, with intraspecific units governed locally by demographic processes.

**HIGHLIGHTS:**

1. Phylogeographies of vertebrates within the southwestern deserts of North America have been shaped by climatic fluctuations imbedded within broad geomorphic processes.
2. The resulting synergism drives evolutionary processes, such as an expansion of within-species genetic divergence over time. Taxonomic inflation often results (i.e., an increase in recognized taxa due to arbitrary delineations), such as when morphological divergences fail to juxtapose with biogeographic hypotheses.
3. However, isolated groups can be discriminated within-species by mapping genetic variability onto geographic distances. This approach can often diagnose ‘hard’ barriers to dispersal, or alternatively, strong selection acting against hybridization. On the other hand, elevated genetic divergences among groups less-isolated would underscore isolation-by-distance (i.e., an increase in genetic differentiation concomitant with geographic distance).
4. The biogeographic patterns we identified in Tiger Whiptail are largely synonymous with those found in other regional species, particularly given the geomorphic separation of Mohave and Sonoran deserts by the Colorado River, and Sonoran/ Chihuahuan deserts by the Cochise Filter Barrier.
5. Our results for the Tiger Whiptail complex broaden and extend the context within which polytypic species are conserved and managed, particularly those that reflect an incongruence among molecular and morphological standards.

## INTRODUCTION

Whiptail lizards (*Aspidoscelis*; Teiidae) comprise one of the most conspicuous groups of lizards in western North America. Their complex history of hybridization/ introgression coupled with multiple transitions to parthenogenic life histories fuels their extraordinary capacity as an evolutionary model (Barley et al. 2019). However, intraspecific morphological variation coupled with high rates of hybridization has made species-delineation difficult, which in turn has muddied their taxonomy. The occurrence of paraphyly in the group was partially resolved by subsequently partitioning species between a resurrected *Aspidoscelis* and the nominate *Cnemidophorus* (Reeder et al., 2002). While a positive step, it also necessitated reallocation of several groups, such as: *Aspidoscelis cozumelus* (parthenogenetic species), *A. deppi* (gonochoristic species), *A. sexlineatus* (parthenogenetic and gonochoristic species), *A. tesselatus* (parthenogenetic species), and *A. tigris* (gonochoristic species). As a result, many systematic issues inherent within *Cnemidophorus* were simply transferred to *Aspidoscelis*.

The *A. tigris* species-group, for example, represents a perplexing systematic challenge as it comprises a widely distributed complex of gonochoristic populations located within desert regions of southwestern North America, northern México (to include Baja California), as well as numerous islands east and west of the peninsula. Although the group *per se* is characterized by a readily assessed set of morphological characters (Walker and Maslin, 1981) and a distinctive karyotype (Lowe et al., 1970), its diversity (i.e., ranging from a single species to more than a dozen) has been a contentious issue in North American Herpetology. Much of its diversity is represented by distinct populations located on relatively small, vulnerable islands in the Gulf of California. Several of these were designated as distinct species (Walker et al., 1966a, b; Walker and Taylor, 1968; Walker and Maslin, 1981), based snout-vent length, color pattern, and/or morphology (e.g., *A. bacatus* of Isla San Pedro Nolasco; *A. catalinensis* of Isla Santa Catalina; *A. martyris* of Isla San Pedro Martír; and *A. celeripes* of Isla San Jose). However, some authors (e.g. Lowe et al., 1970; Wright, 1993) have considered them instead as subspecific variants. Continental populations have also proven taxonomically contentious. For example, a longstanding disagreement (Hendricks and Dixon, 1986; Dessauer et al., 2000) centers on whether *A. tigris* and *A. marmoratus* in small areas in Hidalgo County NM represent intergrading subspecies or hybridizing species.

Taxonomic difficulties are thus quite apparent within the *A. tigris* complex, and primarily stem from complex patterns of intergradation and hybridization confounded by nondiscriminating morphological variation. To provide clarification, we evaluated a complex of four continental subspecies (*A. t. mundus*, *A. t. punctilinealis*, *A. t. septentrionalis*, and *A. t. tigris*; Fig. 1), plus a recently elevated putative sister species, *A. marmoratus* (Hendricks and Dickson, 1986; Reeder et al., 2002; Tucker et al., 2016). Our goal was to examine how mitochondrial (mt)DNA variation in *A. tigris* subspecies compares with morphological discordance [e.g., subspecies seemingly supported by color pattern yet weakly differentiated by scalation and morphometrics (Taylor et al., 1994; Walker et al., 2015; Fig. 2)]. Furthermore, morphological intergrades and overlapping genetic polymorphisms suggest admixture among several groups: *A. t. tigris* and *A. t. septentrionalis* (Taylor 1988), *A. t. tigris* and *A. t. punctilinealis* (Taylor 1990), and *A. t. punctilinealis* and *A. t. septentrionalis* (Dessauer et al. 2000).

**Figure 1:**
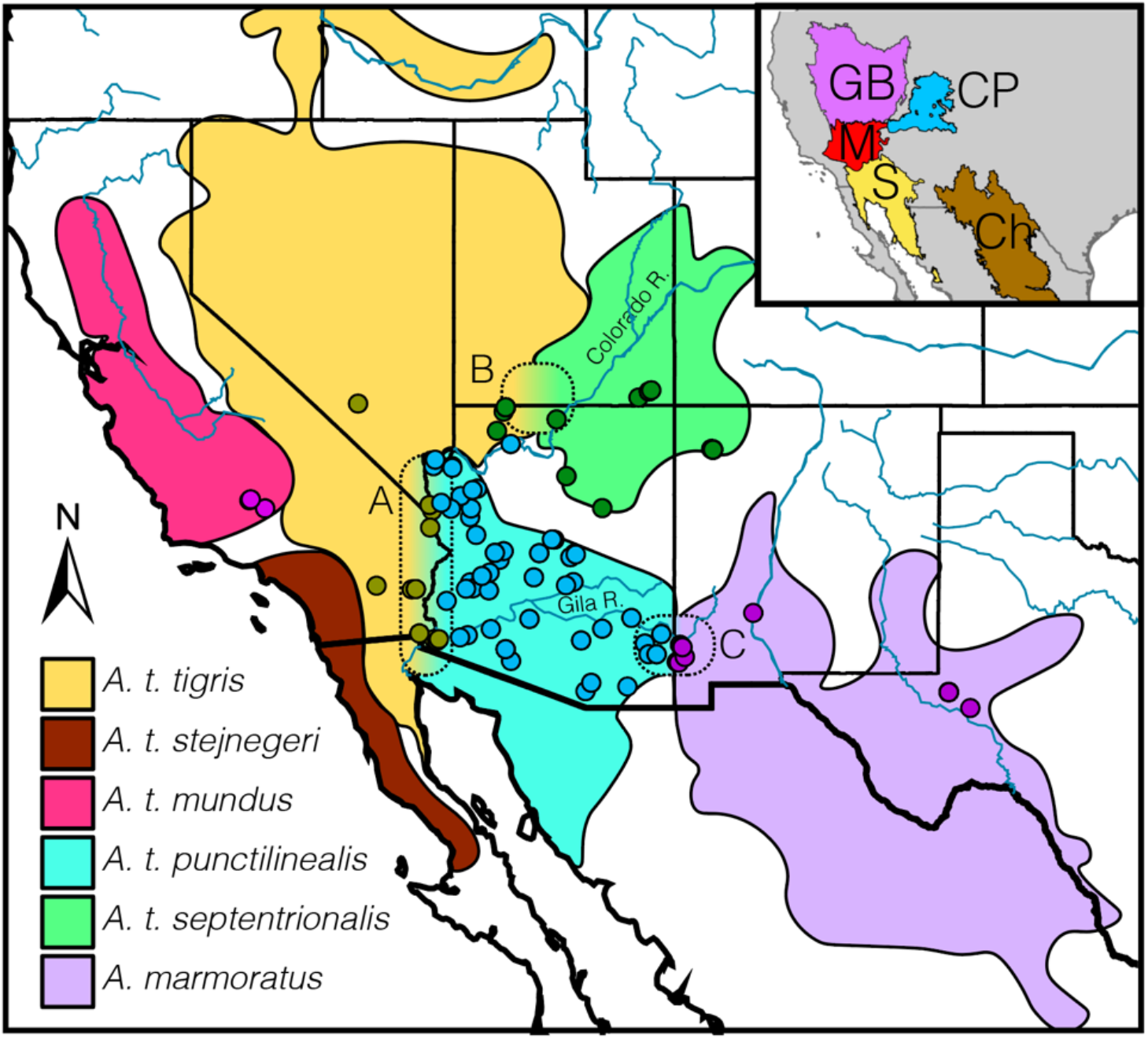
Range map and sampling sites for *Aspidoscelis marmoratus* and subspecies of *A. tigris* (subspecific units within *A. marmoratus* and Mexican subspecies of *A. tigris* not shown). Three identified contact zones are depicted between: (A) *A. t. tigris* and *A. t. punctilinealis*, roughly transected by the Colorado River with morphoclines varying from 16km in the north to ~180km in the south; (B) *A. t. tigris* x *A. t. septentrionalis*, where color-pattern intergradation corresponds to a ~50km transition from Mohave (inset; M) to Great Basin (GB) desert-scrub communities to the Colorado Plateau ecoregion (CP); and (C) *A. t. punctilinealis* x *A. marmoratus*, a hybrid zone of <2km in width traversing the Cochise Filter Barrier transition zone between the Sonoran (inset; S) and Chihuahuan (Ch) deserts. Note that both sample points and ranges are colored by subspecies, as defined in the legend. Map based on observations and prior work by BKS.

**Figure 2:**
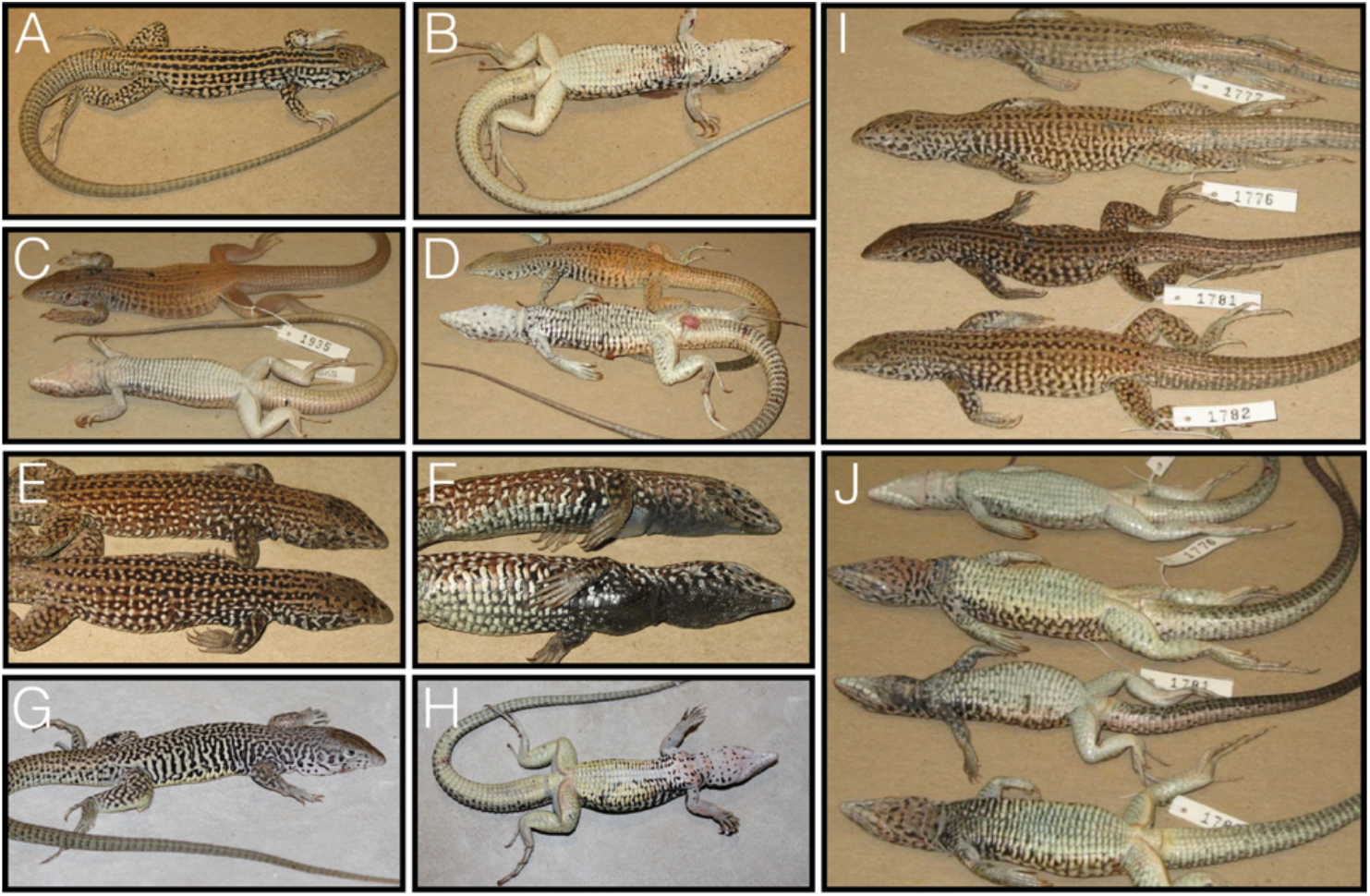
Pattern polymorphism in the *Aspidoscelis tigris* group: (A–B) *Aspidoscelis tigris mundus;* (C) *A. t. septentrionalis* from northern Arizona; (D) *A. t. tigris* from California; (E–F) *A. t. punctilinealis* from central Arizona; (G–H) *A. marmoratus* from New Mexico; (I) Dorsal and (J) ventral view of suspected *A. t. tigris* x *A. t. punctilinealis* hybrids. Pictures provided by BKS.

The contact zone separating *A. t. punctilinealis* and *A. marmoratus* (i.e., *A. t. marmoratus* of the following authors) coincides with a genetic cline at multiple loci (Dessauer and Cole 1991, Dessauer et al. 2000), suggesting secondary contact following a period of isolation (Barton and Hewitt, 1985). A steep cline involving six color-pattern characteristics is also apparent in the hybrid zone between *A. t. punctilinealis* and *A. marmoratus* (Dessauer et al., 2000), arguing for the utility of these characters for delineation of *A. tigris* lineages.

In the present study, we focused on defining phylogeographic patterns and interpreting phylogenetic delineations primarily within the range of *A. t. punctilinealis* in Arizona. To the east, it forms steep morphological/ genetic clines at a hybrid zone with *A. marmoratus* (Dessauer et al., 2000), whereas to the west it exhibits a relatively gradual phenotypic gradient with *A. t. tigris* (Taylor 1990). Interestingly, both *A. t. septentrionalis* and *A. t. punctilinealis* also form exceptionally gradual morphoclines with *A. t. tigris* that lack apparent ecological underpinnings (Taylor et al. 1994, Taylor 1990). This stands in contrast to that found between *A. t. punctilinealis* and *A. marmoratus* (<2km; Dessauer et al., 2000; Zweifel 1962). Given this discontinuity, we also examined the association between morphological gradation in the complex and its genetic discontinuity. To do so, we extended the phylogenetic resolution of lineages within the delineated *A. tigris* complex, then contrasted our phylogenetic patterns with morphological species identifications. We generated taxonomic hypotheses by employing ‘species delimitation’ algorithms, then employed subsequent downstream analyses to compare the manner by which each model partitioned genetic variation.

### The Colorado Plateau as a biogeographic entity

The Colorado Plateau is a tectonically intact landform (~2000m elevation) with scant internal deformation, thus providing a relatively linear configuration of layered sedimentary rock for the Plateau interior that contrasts with an eroding margin since Late Miocene (Levander et al., 2011). It also stands in sharp contrast with the more fractured geomorphic features of neighboring regions: The Rocky Mountains (immediately north and east), Rio Grande Uplift (southeast), and the Basin and Range Province (immediately west) (Fig. 1 of Flowers, 2010). The uplift of the Plateau and the integration of the Colorado River were defining geomorphic events (Minckley et al., 1986; Spencer et al., 2008).

Prior to integration, the Colorado River emptied into an endorheic basin, most likely following the early path of a major modern tributary (Little Colorado River; House et al., 2008). In late Miocene-early Pliocene, headwater erosion and basin ‘spillover’ forged a connection between the river and the Gulf of California via the Salton Trough (Dorsey et al., 2018; Nicholson et al., 2019; Sarna-Wojcicki et al., 2011). The Gila River system, which had previously drained directly into the Gulf (Eberly & Stanley, 1978) was similarly incorporated. Other drainages that terminated into closed basins during the Miocene (i.e. Salt and Verde rivers) were also integrated with the Gila River system by late Pliocene (Blakely & Ranney, 2008). The Pliocene also brought decreased seasonality as well as elevated temperatures and humidity, a climatic regime subsequently reversed by Quaternary glaciation (Thompson, 1991).

Geomorphic events were not only pivotal with regard to biological diversification on the Plateau (Bangs et al., 2020a, 2020b; Douglas et al., 1999; Douglas et al., 2016) but also provoked endemism as well, in synergy with widespread physiographic and climatic oscillations. These created ‘boom-and-bust’ demographic cycles as Plateau habitats expanded and contracted (Douglas et al., 2006), with bookmarks subsequently deposited within the genomes of desert herpetofauna (Douglas et al., 2006; Douglas et al., 2016; Mussmann et al., 2020; Sullivan et al., 2014). Ancillary genetic footprints of hybridization are also increasingly apparent in modern evaluations (Chafin et al., 2020; Bangs et al., 2020a; Bangs et al., 2020b). Opportunities for genetic exchange (or alternatively, vicariant events) hinged largely upon drainage evolution (e.g., Chafin et al., 2020; Douglas et al., 1999). Small-bodied terrestrial vertebrates were also involved in these biogeographic opportunities, despite the fact that the Colorado River had long been hypothesized as a vicariant barrier to dispersal (Lamb et al., 1992; Pounds and Jackson, 1981).

## MATERIALS AND METHODS

### Sampling, amplifying and sequencing of mtDNA

We evaluated a total of 239 samples representing five members of the *A. tigris* complex (*A. marmoratus*, *A. tigris mundus*, *A. t. punctilinealis*, *A. t. septentrionalis*, and *A. t. tigris*), their putative secondary intergrades (e.g., hybrids), plus two out-groups (*Tupinambis rufus* and *Aspidoscelis gularis*) (Fig. 1). Genomic DNA was isolated from tail clippings or liver tissue using the Chelex Instagene-matrix extraction protocol (Bio-Rad).

We utilized published primers and PCR protocols (Douglas et al., 2002) to amplify two overlapping mtDNA genes (ATP8 and ATP6). Poor amplification in *A. marmoratus* also forced the design of species-specific primers (Primer3; Untergasser et al., 2012). These were: ATP_F (3-CGCACCCTGATTTATTGTTT-5) and ATP_R (3-GGGTGAAAACGTATGCTTGA-5). Each 20 μl PCR reaction consisted of 1.5 μl of 1/10x DNA template, 10 μl of HotStarTaq Master Mix (Qiagen), 0.08 μM of each primer, with the reaction volume finalized with distilled water. After verification via agarose gel electrophoresis, amplicons were purified using an Exonuclease I/Shrimp Alkaline Phosphatase. Forward and reverse sequencing reactions were performed using BigDye v3.1 Dye Terminator chemistry (Applied Biosystems, Inc.) and sequenced at the University of Illinois Roy J. Carver Biotechnology Center. Raw sequence chromatograms were manually curated (Sequencher ver. 5.0, Gene Codes, Ann Arbor, MI) and aligned (Mega_5; Tamura et al., 2011).

### Phylogenetic inference

To develop initial hypotheses, we first inferred phylogenetic relationships using both Bayesian and distance-based methods. We tested models of nucleotide substitution (N=88-JModelTest; Darriba et al., 2012), using Akaike and Bayesian information criteria (AIC and BIC, respectively). We generated an ultrametric tree using the HKY-model (selected by both AIC and BIC) with gamma distributed rates and invariant sites (BEAST2; Bouckaert et al., 2014). We applied a Yule model with default parameters and a relaxed log-normal clock process. MCMC chains were sampled every 500^th^ of 10 million MCMC iterations, following a burn-in period of 2 million. Chain convergence and mixing were visually assessed, and MCMC samples thinned so as to maintain effective sample sizes >200 while reducing autocorrelation of samples (Tracer; Rambaut and Drummond, 2007). The posterior sampled trees were summarized as a maximum clade credibility tree with branch lengths representing median heights. For contrast, we also generated a neighbor-joining (NJ) tree using model-corrected distances (R-package ape; Paradis et al., 2004) in R (R Core Team, 2020), with nodal support assessed via 100 bootstrap pseudo-replicates, as well as a maximum-likelihood tree in PHYML (Guindon et al. 2010). Basic population summary statistics for major phylogenetic clades (as diagnosed via nodal support and statistical tests of monophyly) were also computed in PEGAS (Paradis, 2010).

### Delimiting groups within A. tigris

We first evaluated the currently accepted taxonomic designations for subspecies within *A. tigris* (e.g., Crother et al., 2017; hereby abbreviated as ‘TAX’), then employed an array of analytical approaches to generate six additional delimitation hypotheses. Distance-based clustering methods, used in the context of species delimitation, assume that biological units (i.e., species) form non-overlapping genetic clusters. We first employed a recursive method for ‘Automated Barcode Gap Discovery’ (ABGD; Puillandre et al., 2012) that statistically infers a threshold distance from the data (i.e., the genetic distance at which groups are considered to represent different clusters). We also generated clusters using corrected genetic distances as input (CD-HIT; Fu et al., 2012), with manually specified distance thresholds of 2% and 5% (hereafter referred to as CD2 and CD5). These hard thresholds were chosen because they represent approximate ‘breakpoints’ at which changes occur in the ABGD analysis regarding the number of sampled groups.

We defined phylogenetic species using several approaches: The first, a ‘naïve’ method, defined groups as the shallowest divergences statistically supported and reciprocally monophyletic (Rosenberg, 2007), hereafter abbreviated as ‘ROS’ (Spider R package; Brown et al., 2012). A second approach derived groups from tree characteristics under an assumed speciation process. The first of these methods [‘Generalized Mixed Yule Coalescent’ (GMYC; Pons et al., 2006)] represents the species boundary as a distinguishable change in branching patterns within the phylogeny, with independently evolving clusters diagnosed as those at which branching patterns transition from expected under two different models: An assumed interspecific (Yule process; Nee et al., 1994) and an intraspecific (neutral coalescent; Hudson, 1990). In the second [‘Poisson Tree Process’ (PTP; Zhang et al., 2013)] the probability of speciation is considered in terms of mutational distance between cladogenesis events (as assumed with a Poisson distribution). PTP requires a tree with branch lengths scaled per substitutions, and given this, our input represented a tree topologically consistent with that inferred by BEAST but derived instead with PHYML (Guindon et al. 2010). GMYC was run using the Splits R package (Fujisawa et al., 2013); PTP using the Bayesian implementation provided by the bPTP server (Zhang et al., 2013).

### Comparingf delimitations

We used 2-way and 3-way Analyses of Molecular Variance (AMOVA; Excoffier et al. 1992) with group assignments defined by individual membership under each hypothesis. We then used isolation-by-distance (IBD) as a null model under the assumption that the entire complex represented a single species. Our justification stemmed from the pervasive nature of IBD with regard to dispersal-limited species (Sexton et al. 2014). In this sense, deviations from the expected IBD pattern are hypothesized as diagnostic of some alternative non-spatial process (e.g. reproductive isolation). Partial Mantel tests (Smouse et al. 1986), employing pairwise genetic distances with geographic distances as a covariate (Douglas and Endler, 1982; Meirmans, 2012), tested the fit of individual assignments under each delimitation hypothesis. Simple and partial Mantel tests (R-package vegan; Oksanen et al., 2007) utilized corrected genetic distance (ape; Paradis et al., 2004) with confidence assessed using 9,999 permutations. We also employed a stratified Mantel test (per Meirmans, 2012) with permutations constrained within clusters, to test if significance of the IBD model is impacted by sampling across structured populations, as delimited by each of our hypotheses.

We also used a spatially explicit partitioning analysis with genetic variance [measured as a fixation index (*F*_CT_)] maximized among groups by iteratively rearranging samples into *K*-clusters [Spatial Analysis of Molecular Variance (SAMOVA); Dupanloup et al., 2008]. These were executed for *K*-values 2-20, using 100 bootstrapped alignments and replicated without explicit geographic constraints. We also identified spatial transitions in genetic distances using an interpolation approach [Alleles-in-Space (AIS); Miller, 2005].

## RESULTS

### *Phylogenetic and spatial relationships within* Aspidoscelis tigris

The final 777bp alignment resulted in 120 haplotypes. Bayesian analysis (BEAST2) yielded strong support for *A. t. tigris* + *A. t. punctilinealis* (ROS clades A–F, Fig. 3; also see Supplementary Figure S1) as being sister to *A. t. mundus* + *A. t. septentrionalis* (ROS clade G). All clades were significantly supported as being reciprocally monophyletic [Rosenberg (2007) test], and all reflected similarly high bootstrap support in the neighbor-joining analysis. The maximum-likelihood analysis resulted in a tree topologically identical, and was thus not displayed.

**Figure 3:**
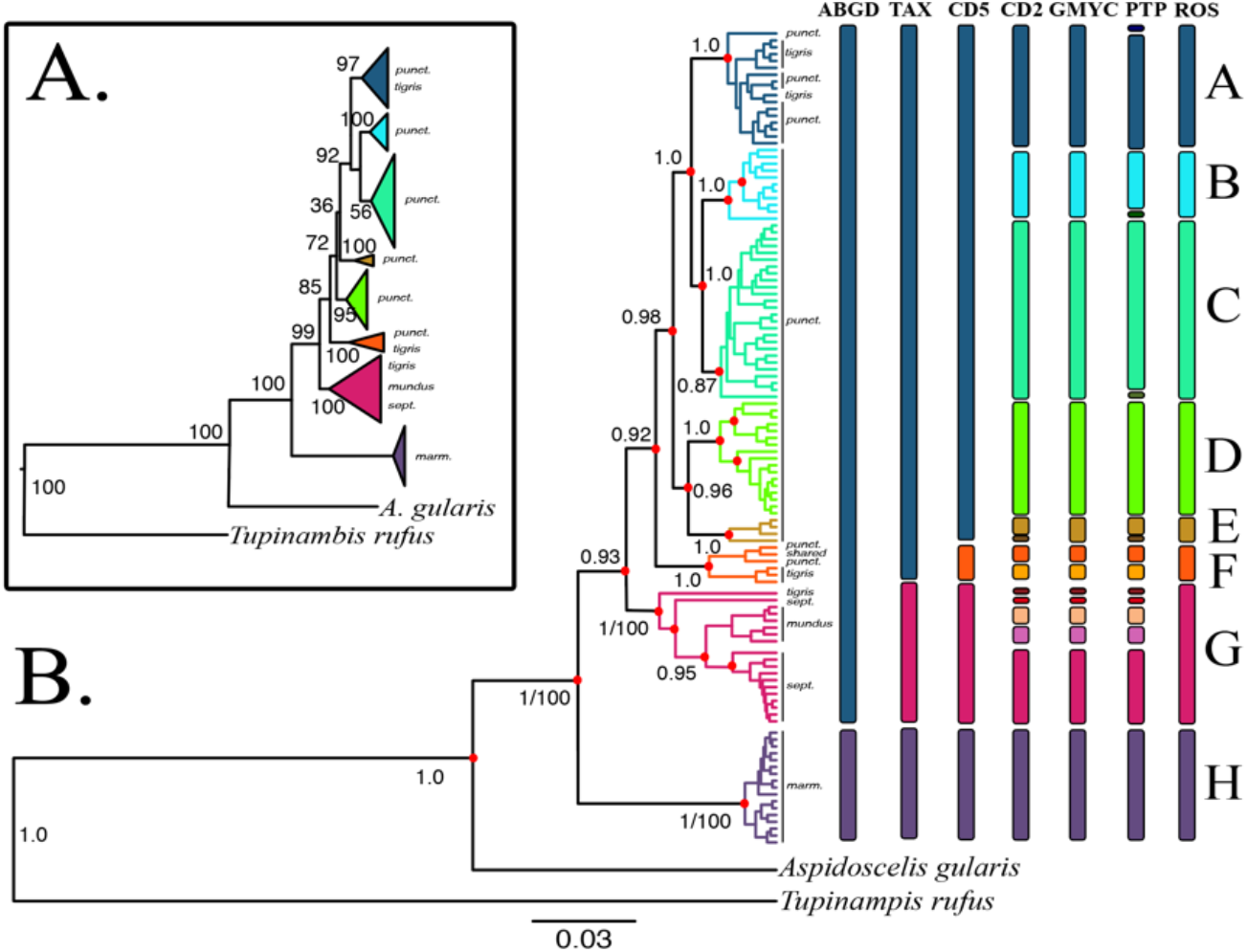
Neighbor-joining (A) and Bayesian (B) phylogenetic trees based on sequence analysis of 777bp of the ATPase 6 and 8 genes across 239 samples, contrasted against results of *de novo* species delimitation models represented by colored bars. Branches are labelled according to the morphological diagnoses of the tissue samples from which mtDNA sequences originated and are abbreviated as follows: *tigris=A. tigris tigris;punct.=A. t. punctilinealis; sept.=A. t. septentrionalis; mundus=A. t. mundus; marm.=A. marmoratus; shared=found* in *A. t. tigris* and *A. t. punctilinealis*. Model abbreviations are: ABGD=Automated Barcode-Gap Discovery; TAX=qualitative hypothesis derived from current taxonomy; CD5=CD-Hit with 5% divergence threshold for clustering; CD2= CD-Hit with 2% divergence threshold for clustering; GMYC=Generalized mixed Yule coalescent; PTP=Poisson tree process; ROS=Clade delimitation based on statistical tests of monophyly. Node labels represent posterior probabilities from Bayesian inference (B), and neighbor-joining bootstrap support (A; omitted for shallow nodes). Nodes passing the statistical test of reciprocal monophyly are indicated with a red dot. Clades in both trees are colored according to the ROS model.

All four subspecies formed a monophyletic group sister to a well-defined *A. marmoratus* (clade H). *Aspidoscelis t. septentrionalis* (clade G, excluding *mundus* samples) demonstrates significantly negative values for both Tajima’s and Fu and Li’s *D* statistics, with exceptionally low nucleotide diversity and haplotype diversity much less than that recorded for *A. t. punctilinealis* and *A. t. tigris* (Table 1). These suggest a population expansion following a bottleneck (Douglas et al., 2010). *Aspidoscelis marmoratus* showed similar signatures, although with a much greater differential between haplotype and nucleotide diversities, and a reduced number of segregating sites relative to sample size (Table 1). Together, *A. t. tigris* + *A. t. punctilinealis* were subdivided into five major clades with strong nodal support and reciprocal monophyly (clades A–F, Fig. 3).

**Table 1:** Population summary statistics for clades within the *Aspidoscelis tigris* complex as determined by sequence analysis of 777bp of the ATPase 6 and 8 genes across 239 samples. Abbreviations are as follows: *H* = haplotype diversity; *π* = nucleotide diversity; *θ* = population mutation rate; *θ_s_/N* = number of segregating sites / clade sample size; *H/N* = number of haplotypes / clade sample size. *A. t. mundus* was excluded due to low sample size.

| CLADE | TAJIMA<br>*D* | TAJIMA *D*<br>P-VALUE | *H* | $\pi$ | $\theta$ | FU & LI<br>*F* | FU & LI<br>*D* | $\theta_s/N$ | *H/N* |
| --- | --- | --- | --- | --- | --- | --- | --- | --- | --- |
| A | -1.522 | 0.128 | 0.993 | 0.013 | 15.409 | -1.680 | -1.595 | 2.889 | 0.944 |
| B | -0.938 | 0.348 | 0.969 | 0.011 | 11.258 | -0.442 | -0.356 | 2.833 | 0.916 |
| C | -1.628 | 0.103 | 0.997 | 0.012 | 17.033 | -2.051 | -1.935 | 2.461 | 0.923 |
| D | -1.889 | 0.059 | 0.966 | 0.018 | 26.468 | -2.968 | -3.008 | 3.885 | 0.654 |
| E | -0.524 | 0.600 | 1.000 | 0.013 | 10.363 | 0.050 | 0.524 | 4.75 | 1.000 |
| F | 0.616 | 0.538 | 1.000 | 0.021 | 14.890 | 0.660 | 0.374 | 5.667 | 1.000 |
| G | -2.114 | 0.017 | 0.478 | 0.006 | 15.600 | -2.609 | -2.123 | 1.123 | 0.292 |
| H | -1.778 | 0.075 | 0.828 | 0.002 | 3.762 | 0.457 | -2.933 | 0.4 | 0.425 |

Morphologically-diagnosed *A. t. tigris* was polyphyletic, with all samples but one allocated to either clade A or F. The single exception was a morphologically diagnosed *A. t. tigris* (Nevada) that fell within the *A. t. septentrionalis/mundus* haplogroup (clade G). Clade F, which was >2% divergent from the remaining *A. t. tigris/punctilinealis* clades (A–E), contained both morphologically consistent *A. t. tigris* (Clark County, southern Nevada), as well as a small subset of *A. t. punctilinealis* (Mohave County, northwest Arizona). All other morphologically identified *A. t. tigris* from California had mitochondrial haplotypes aligning with clade A of *A. t. punctilinealis*. This clade formed a monophyletic grouping in the BA analysis with samples from southeastern Arizona (clades D and E). Spatially, clades D+E are separated by the mainstem Gila River and defined to the east by the Cochise Filter Barrier that separates Sonoran and Chihuahuan deserts (Castoe et al. 2007).

All clades within *A. t. tigris/punctilinealis* generally showed higher numbers of segregating sites relative to nucleotide diversity (negative values for Tajima’s *D*; clades A–E Table 1). Clades A-D also showed a high rate of singleton haplotypes. Although non-significant (Table 1), they suggest past population expansions. Within the remainder of the *A. t. punctiliealis*, clade A (southwestern Arizona) is sister to a monophyletic group composed of clade B (Grand Canyon and Colorado River mainstem at juncture between Arizona, Nevada, California), and clade C (central Basin and Range physiographic region of Arizona). The latter is bounded on the north by the Mogollon Rim and is separated from clade B to the south by the Gila River.

The appearance of clade C haplotypes south of the Gila River mainstem indicates limited gene flow between the two groups, possibly reflecting trans-riverine dispersal as promoted by low seasonal flows. Similarly, clade B haplotypes are found east and west of the Colorado River mainstem, again suggesting a riverine barrier historically permeable to dispersal, or with mitochondrial exchange facilitated by shifts in flow. However, no apparent geographic features support the phylogenetic separation of clades A and C. Their sister relationship is also sustained in the NJ analysis, although with a bootstrap support of 56% (compared to 0.87 posterior probability in the Bayesian phylogeny).

### Species delimitation models

We found considerable variation in the number of proposed clades generated by species delimitation models. Fig. 3B depicts the manner by which each model maps onto the phylogeny, using an identification scheme consistent with Fig. 4 and Supplementary Figure S1. Phylogenetically defined clades (‘ROS’) and a taxonomic hypothesis (‘TAX’) both collapsed *A. t. tigris* and *A. t. punctilinealis* on the basis of extensive intergradation (Taylor et al. 1994).

**Figure 4:**
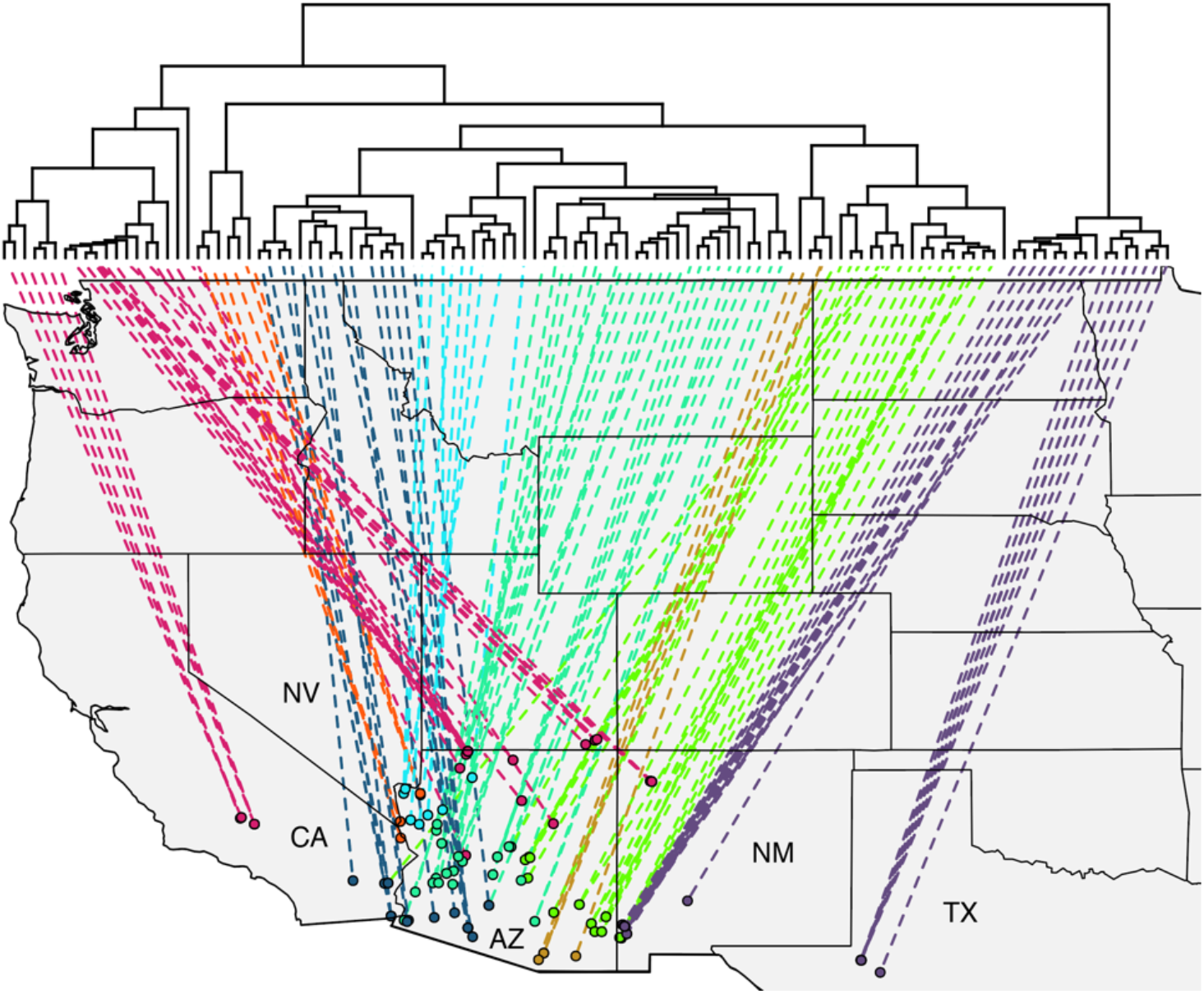
Geo-referenced haplotype tree generated via Bayesian analysis, representing 777bp of the ATPase 8 and 6 genes across 239 samples. Clade coloring follows the ROS model (see Fig. 1)

The ABGD analysis was by far the most conservative, collapsing both *A. t. punctilinealis* and *A. t. tigris* with *A. t. septentrionalis* + *A. t. mundus*, while only segregating *A. marmoratus* (Fig. 3B). CD-HIT, with an *a priori* sequence divergence threshold of 5%, produced a 4-species model (CD5) that separated *A. t. septentrionalis* + *A. t. mundus, A. marmoratus*, and *A. t. punctilinealis + A. t. tigris*. The latter was also partitioned into two groups (i.e., separating clade F). One consisted of haplotypes from northwestern Arizona and southern Nevada (clade F; a subset of most morphologically consistent *A. t. tigris*), while the second comprised *A. t. punctilinealis* (clades A–E) and remaining *A. t. tigris*, to include those with ranges overlapping clade F. Decreasing the threshold to 2% (i.e., model CD2) then divided *A. t. punctilinealis* into eight groups, and five representing *A. t. septentrionalis* (i.e., two from California and three occupying the Colorado Plateau; Fig. 4). GMYC and PTP produced large numbers of groups, with 13 in the former and 16 in the latter, with many consisting of but a single haplotype.

### Partitioning of genetic variance

The spatial analysis supported models with fewer species. In the bootstrapped datasets, *F*_CT_ overlapped among *K* values (Fig. 5), with the greatest increase in among-group variance (Δ*F*_CT_) occurring at *K*=3. Individual assignment at *K*=2 corresponded to the ABGD model (two species: *A. marmoratus* and *A. tigris)*, with three groupings in *K*=3: *A. marmoratus; A. t. septentrionalis* + *A. t. mundus*; and *A. t. punctilinealis* + *A. t. tigris*.

**Figure 5:**
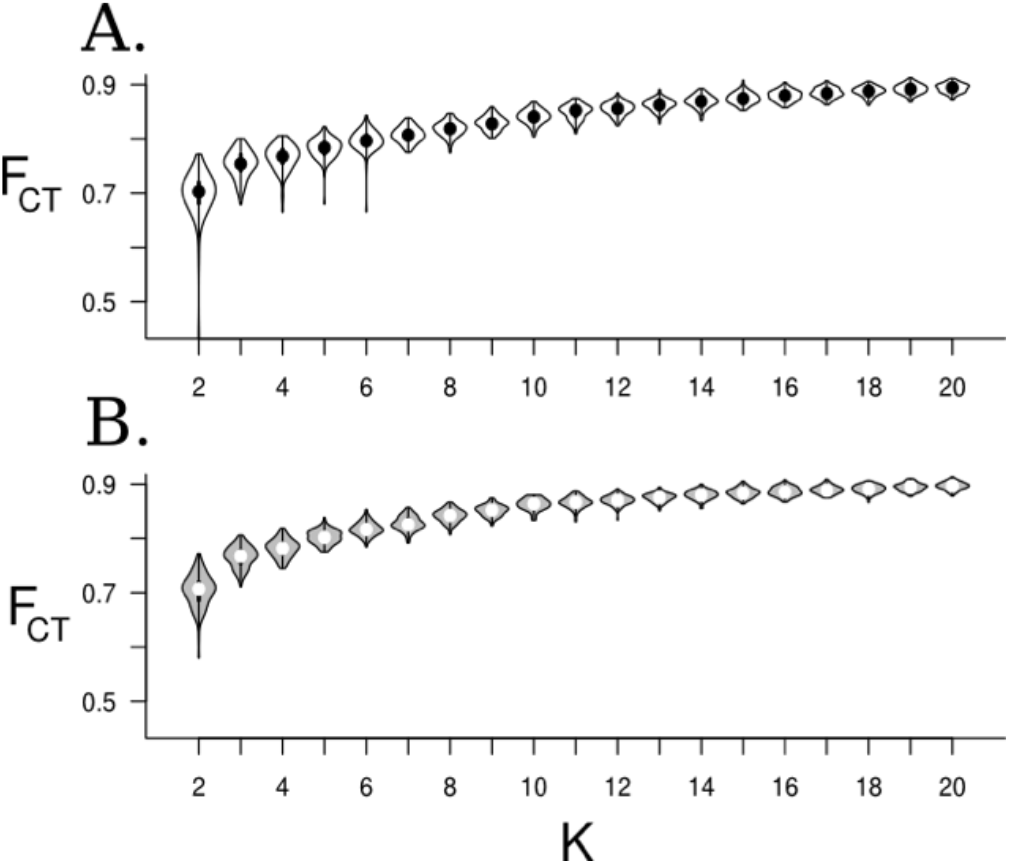
Among-group genetic variance (*F*_CT_) for genetic clusters within the *Aspidoscelis tigris* complex, as derived by SAMOVA clustering runs. Cluster optimization constrained by (A) geographic proximity of samples and (B) without a geographic constraint. The x-axis (*K*) reports the number of genetic clusters sampled.

A significant correlation was found between matrices of pairwise genetic and geographic distances (Mantel test: r=0.576, P=0.0001; Table 2; Fig. 6). Significant associations only occurred with the ROS, TAX, ABGD and CD5 models when permutations were limited to those strata defined by delimitation models (i.e., stratified Mantel). The partial Mantel tests with the ABGD model produced the strongest association (r=0.777), with values generally declining as the number of species increased (Table 2). Correlation of group membership with genetic distance under all models was greater than in the simple Mantel test for all cases, save for the PTP model (r=0.5754), suggesting that all delimitations better encapsulate spatial genetic variation than one in which all examined samples are collapsed.

**Figure 6:**
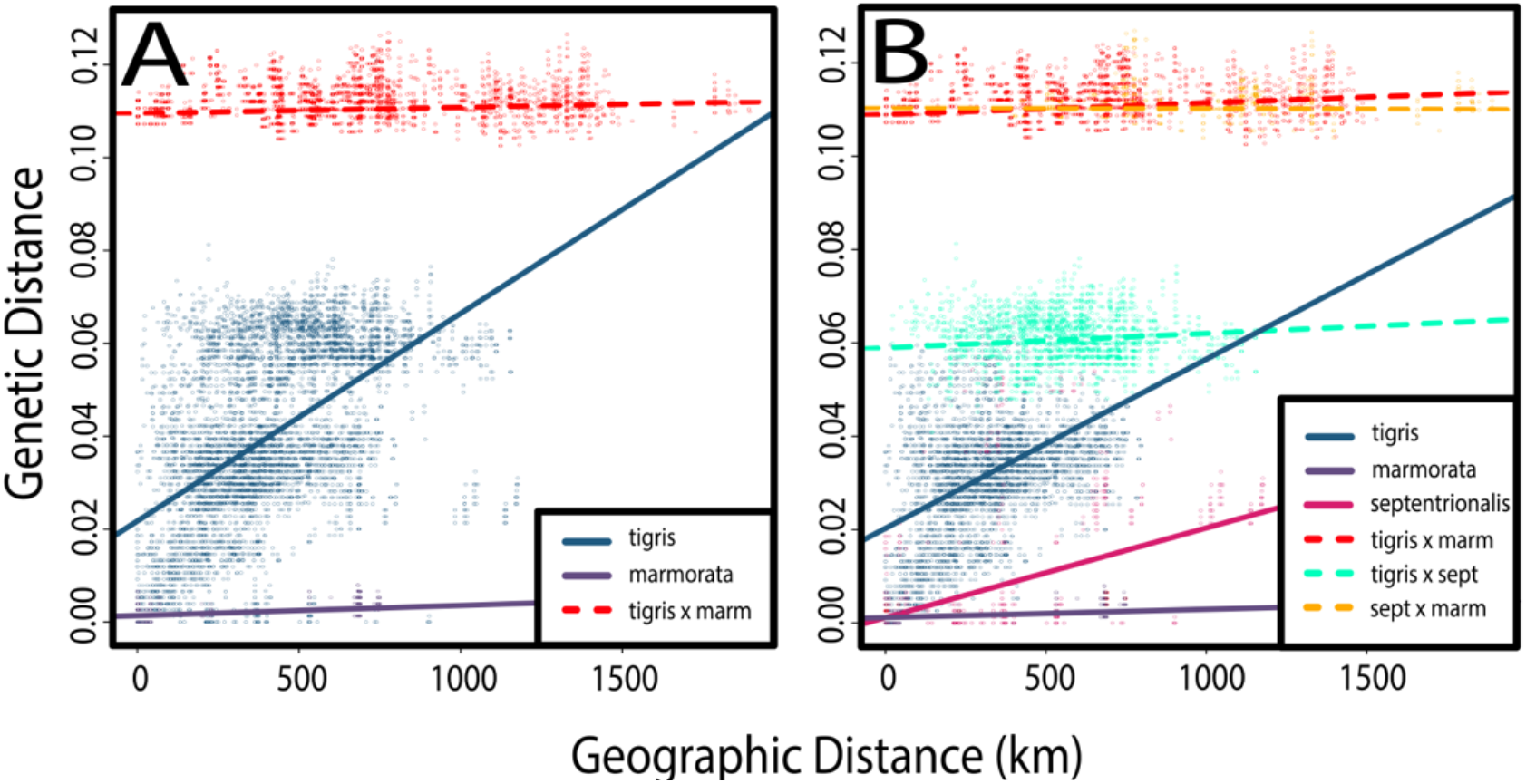
Within- and among-group linear regression of geographic x genetic distances for clades delimited by (A) the ABGD (Automated barcode-gap discovery) model and (B) TAX (taxonomic) model. Coefficients of correlation (*r*^2^) ranged from 0.01-0.449. Note that “*tigris*” refers to both *A. t. tigris* and *A. t. punctilinealis*, as these subspecies are not differentiated in the models. *A. t. mundus* was excluded due to the geographically restricted sampling presented herein.

**Table 2:** Comparison of alternative species delimitation models according to the fit of the data under an isolation-by-distance model. Results are shown for 2-way, stratified, and partial Mantel tests, reporting a correlation coefficient (*r*) between distance matrices composed of geographic and genetic distances. Correlation coefficients in the partial Mantel tests reflect association between clade membership and genetic distance, with geographic distances as a covariate. Values indicated by (**) are significant at p>0.0001. Species delimitation models are abbreviated as: ABGD=Automated Barcode-Gap Discovery; TAX=qualitative hypothesis derived from current taxonomy; CD5=CD-Hit with 5% divergence threshold for clustering; CD2= CD-Hit with 2% divergence threshold for clustering; GMYC=Generalized mixed Yule coalescent; PTP=Poisson tree process; ROS=Clade delimitation based on Rosenberg test of reciprocal monophyly.

| MODEL | 2-WAY<br>MANTEL TEST | STRATIFIED 2-WAY MANTEL | PARTIAL<br>MANTEL |
| --- | --- | --- | --- |
| | Mantel $r$ | Mantel $r$ | Mantel $r$ |
| ALL | 0.5762** | - | - |
| ROS | - | 0.5762** | 0.6317** |
| ABGD | - | 0.5762** | **0.7770\*\*** |
| TAX | - | 0.5762** | 0.7118** |
| CD5 | - | 0.5762** | 0.7025** |
| CD2 | - | 0.5762 | 0.5816** |
| GMYC | - | 0.5762 | 0.5818** |
| PTP | - | 0.5762 | 0.5754** |

\*\*Significant at $\alpha=0.0001$

Overall, those models partitioning *A. tigris* subspecies were supported by AMOVA analyses (Supplementary Table S1, S2), with high genetic variance among groups rejecting panmixia. Among-group genetic variance (analogous to *F*_CT_) varied between 72–90% and was maximized in models that recognizing greater numbers of partitions (i.e., CD2, GMYC, and PTP), with 89-90% of variance explained. These values generally declined as models became more conservative.

Interpolating the AIS results with a diffusion model (Fig. 7) produced apparent peaks in relative pairwise genetic diversity at contact zones between *A. t. punctilinealis* (in this case clade D) and *A. marmoratus* (clade H). These occurred in southeastern Arizona near the shared border with New Mexico, and in northwestern Arizona, near the contact zone with *A. t. tigris* clade F.

**Figure 7:**
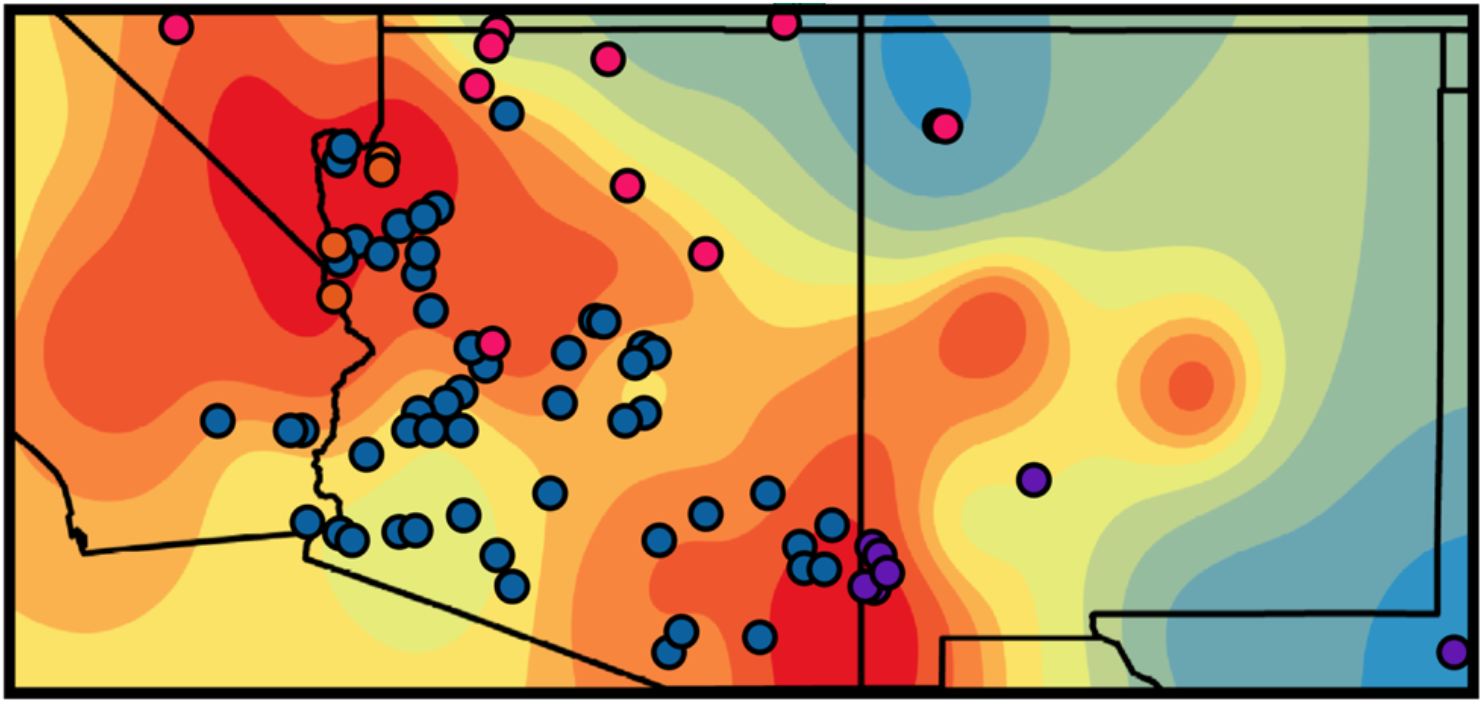
Map depicting results of an ‘Alleles in Space’ analysis for members of the *Aspidoscelis tigris* complex. Relative genetic differentiation is interpolated across the landscape, with individuals (as dots) colored by the CD5 model (see Fig. 1B). Heat map colors depict relative genetic distances per unit of geographic distance, with red corresponding to a greater genetic distance, and blue being lesser genetic distance.

## DISCUSSION

We used mtDNA sequence data to: Infer phylogenetic and phylogeographic patterns within the *Aspidoscelis tigris* complex; Generate species delimitation hypotheses therein; and compare model fit using several approaches. We judged how well each model partitioned genetic variance, both hierarchically and spatially. Our results supported *A. marmoratus* as a distinct species. However, our weak phylogenetic recognition of *A. t. tigris* and *A. t. punctilinealis*, as well as *A. t. mundus* and *A. t. septentrionalis* questions their subspecific designations, and suggests the need for additional work utilizing larger number of molecular markers to assess potential intergradation. However, pairwise genetic divergences among the *A. t. tigris/punctilinealis* and *A. t. septentrionalis/mundus* groups, regardless of their geographic proximity (e.g., <10km versus >1000km) may indicate reproductive isolation, again prompting further study that incorporates nuclear markers.

### *Biogeography of speciation in* Aspidoscelis tigris

Deep relationships within the *A. tigris* complex generally supported the current taxonomy and reflected divisions among biotic regions. Here, we specifically refer to *A. t. septentrionalis* on the Colorado Plateau, *A. marmoratus* within the Chihuahan Desert, *A. t. punctilinealis* in the Sonoran Desert, and *A. t. tigris* within Mohave and Great Basin deserts. Haplotypes of *A. t. mundus*, restricted to the California Central Valley, were inferred as monophyletic but nested within a larger *A. t. septentrionalis* clade. This highlights the need for increased sampling, particularly for *A. t. mundus* and *A. t. tigris*, whose distributions were sampled less thoroughly.

Topographically, the separation of *A. t. punctilinealis* and *A. marmoratus* aligns with the Cochise Filter Barrier (CFB), a biogeographic entity that has similarly been implicated as influential in defining phylogeographic relationships among other taxa (e.g., Castoe et al., 2007; O’Connell et al., 2017; Morafka, 1977). Pyron et al. (2015) invoked this barrier as a potential driver of so-called ‘soft allopatry’ (Hickerson and Meyer 2008), in which niche differentiation drives speciation *in lieu* of a hard barrier to dispersal. Such barriers can often lead to permeability at contact zones, given that divergence is maintained by adaptation rather than physical separation. As might be expected, a well-characterized hybrid zone separates *A. t. punctilinealis* and *A. marmoratus*, replete with steep clines in color polymorphism (Dessauer et al., 2000). We also identified a distinct east-west divide in mitochondrial haplotypes that is concordant with an hypothesis of soft allopatry.

### The potential for hybridization

We found phylogenetic subdivision less clear among subspecies of *A. tigris* and surmise this likely reflects a cumulative effect of contemporary physiography coupled with historic processes. *Aspidoscelis t. tigris* and *A. t. punctilinealis* were not distinct. Both transverse the Colorado River, and neither is strictly defined by the Mohave-Sonoran transition. This is perhaps not surprising, given that morphological intergradation between *A. t. tigris* and *A. t. punctilinealis* does not follow a strict ecological gradient, but instead manifests itself as an exceptionally wide and smooth morphocline (Taylor et al. 1994).

*Aspidoscelis t. tigris* from the most northerly part of the range had mitochondrial sequences aligning instead with *A. t. septentrionalis*, potentially suggesting exchange between the two (Taylor, 1988). Here, we emphasize that increased genetic sampling is necessary so as to understand the nature of this putative exchange. In contrast, *A. t. punctilinealis* and *A. t. septentrionalis* were demarcated according to recognized physiographic provinces, with *A. t. septentrionalis* occupying the Colorado Plateau and *A. t. punctilinealis* the Sonoran Desert (Fig. 4, S1). This was expected, particularly given our understanding of their respective ranges (e.g. Fig. 1) and associations with the disparate thorn-scrub communities therein (e.g., Zweifel, 1962; Pianka, 1970). We suggest this ecological gradient represents a ‘soft boundary,’ similar to those that mediate the transition between *A. t. punctilinealis* and *A. marmoratus*. Unique communities characterize both the Colorado Plateau and Sonoran Desert (e.g., Douglas et al., 2002, 2010), and this suggests a common, underlying process driving diversification therein (Riddle and Hafner, 2006; Pianka, 1967).

### The validity of taxonomic designations

Genetic delineation of *A. t. tigris* was inconsistent with traditional taxonomy, in that specimens allocated morphologically to *A. t. tigris* were not monophyletic within the phylogenetic analyses, nor were those groupings consistent with the historically accepted range boundaries for *A. t. tigris* and *A. t. punctilinealis*. Our expectations were that haplotypes sampled from the Great Basin and Mohave deserts (i.e., those regions encompassing *A. t. tigris*) would be monophyletic. Instead, we found a closer alignment with *A. t. septentrionalis* to the northeast and *A. t. punctilinealis* to the southeast. This is likely a result of mitochondrial exchange among these subspecies, again highlighting the need for a genomic evaluation.

Taylor et al. (1994) demonstrated the existence of a wide, smooth morphocline >180 km in width that traversed the Colorado River and separated *A. t. tigris* from *A. t. punctilinealis* along the southern extent of their contact (Fig. 1A). This stands in contrast to a 16 km-wide morphocline found at the northern extent of their contact, and that discrepancy begs an obvious ecological explanation (Taylor 1990). The northern morphocline coincides with the Arizona-California-Nevada nexus, a hotspot for mitochondrial diversity as well, where sub-clades overlap without clear physical boundaries (Fig. 4, S1). In addition, multiple clades (A and F) traverse the Colorado River, with the highly divergent clade F (found primarily within morphologically defined *A. t. tigris*) possibly representing a more widespread Mohave clade.

One explanation for the above variation in clinal width evokes an historic perspective wherein the northern contact between *A. t. tigris* and *A. t. punctilinealis* is secondary, whereas range overlap to the south was maintained. This would generate morphoclines of varying ages and would offer one potential explanation for the variance in mitochondrial divergence (i.e., elevated to the north, nonexistent to the south), as well as morphocline width. Here, we posit that the duration and extent of contact may be a driving mechanism. A related hypothesis would be that the width of the cline is in response to an underlying environmental gradient, such as the transition between Sonoran and Mohave habitats. However, neither hypothesis can be tested with the data presented herein.

An additional possibility is the fluctuating capacity of the Colorado River as a barrier to dispersal. We offer scant evidence for it serving as a vicariant structure, a result corroborated by studies of other reptiles (e.g., desert tortoise: Lamb et al., 1989; iguanids: Lamb et al., 1992). Although the Colorado River does seemingly constrain dispersal (Pounds and Jackson 1981), gene flow could still be facilitated by the periodic fluctuations in its trajectory and flow. In fact, the shallow divergences found among clades transecting the river provide evidence for how recent these events were.

### Climate driven population oscillations

One unexpected finding was the exceptional genetic divergences among *A. t. punctilinealis* clades (A–E). One possible explanation evokes fluctuating habitat availability over time, rather than contemporary features of the landscape. Concordance in relative crown ages among the majority of *A. t. punctilinealis* clades (A–E; Fig. 1B) could suggest that intraspecific divergence of these groups shares a common cause. Douglas et al. (2006) offered an applicable model that explained genetic structuring and population expansion in four desert rattlesnake species as a ‘boom-and-bust’ demographic cycle, caused by historical climate fluctuations. Under this model, a ‘refugium phase’ (=‘bust’) condensed continuous distributions into smaller, allopatric populations such that restricted gene flow promoted genetic drift as manifested by reduced effective population size (*N_e_*). As climate shifted, desert habitat expanded such that ecological barriers separating refugia were removed, allowing distributions to expand (=‘boom’).

These oscillating contractions and expansions placed characteristic molecular signatures within populations, most prominently as a loss of genetic diversity and an exaggerated differentiation among sub-populations post-expansion (Hewitt 2000). This is reflected in an increased number of low-frequency haplotypes, and low nucleotide diversity among contemporary haplotypes (Table 1). We also noted similar patterns in *A. t. septentrionalis* and *A. marmoratus*, suggesting bottleneck-expansion events within these taxa as well.

The effects of the ‘refugium phase’ can be seen when examining modern short-range endemics [i.e., those with distributions easily contained within a 100×100 km grid (i.e., <10,000 km^2^]. An example would be the isolated New Mexico Ridge-nosed Rattlesnake (*Crotalus willardi obscurus*) that reflects signals of relatively recent divergence and population reduction (Holycross and Douglas 2007). Its historic connectivity in the Sky-islands has been truncated by contemporary climate change, whereas its refugium phase, as gauged by demographic and niche modeling, underscored a continued loss of habitat (Davis et al. 2015). We suspect a similar refugium-expansion model would explain the extant genetic differentiation found in the present study.

### Comparing species delimitation approaches

We found considerable variance in the number of groups delimited using the various algorithmic methods, especially when contrasted with the current taxonomy (Fig. 3). This could reflect the capacity of the various models to diagnose cryptic lineages, or alternatively, suggest the data violate implicit assumptions (Carstens et al. 2013). Phylogenetic models (i.e., ROS, PTP, GMYC) generally produced the largest numbers of groups, but did not explain the geographic patterns of genetic variation as well as did other models (Table 2; Figure 3). All models, however, captured greater genetic variance among groups (i.e., 72-90% in AMOVA) than within-group (10-28%), indicating the recognition of an authentic biological signal (Table S1, S2).

Species delimitation employing the coalescent, such as GMYC, have previously been criticized for a tendency to over-split when applied to structured populations (Lohse 2009). This, in turn, suggests that a single-threshold delimitation model may be inappropriate for use with hierarchically structured data (Talavera et al. 2013). Strong intraspecific structure in *Aspidoscelis*, similar to that found in other desert reptiles (Holycross and Douglas 2007, Douglas et al. 2002), may explain the over-fitting of GMYC. A more comprehensive test using additional scenarios is warranted, as this will more appropriately characterize those biases that can emerge when the method is applied to empirical data.

We also employed a second phylogenetic method (PTP) that essentially modeled speciation as a clock-like process, with its probability of occurrence represented as a function of time (i.e., branch length). Given this, any aspect that would promote rate heterogeneity could be sufficient to violate the model, such as the extreme demographic fluctuations suspected for *Aspidoscelis* populations.

Clustering-based methods, on the other hand, may be biased by elevated genetic distances or incomplete sampling, with group delimitation being driven entirely by the arbitrary nature of the (user-selected) threshold. The ABGD model attempts to mitigate this by automatically detecting distance-based partitions, as bounded by priors that account for maximum allowable intraspecific divergence (Puillandre et al. 2012). ABGD and clustering methods appeared to delimit groups that more appropriately represented the genetic/ geographic distances paradigm. When examining genetic distances defined by these methods within-versus among-group, we found *A. marmoratus* approximately equal to *A. tigris* with regard to genetic distances, regardless of whether *A. tigris* was intact or partitioned (Fig. 6). Among-group genetic distances were equal regardless of how proximate were the samples (i.e., genetic x geographic distance slope approaches zero). This suggests that divergence is maintained not by physical distance but instead by some intrinsic qualities possessed by those groups, such as reproductive isolation or mutually exclusive habitat requirements (Fig. 6). In this case, near-zero slopes in the among-group comparisons lend some validity to a species-level designation for *A. marmoratus* and suggested the potential for reproductive isolation within the *tigris* complex (i.e., between *A. t. tigris*/*punctilinealis* and *A. t. septentrionalis*; Fig. 6B). However, reduced sample sizes for *A. t. mundus* and *A. t. tigris* remain an issue. Within-group comparisons resulted in a positive relationship with geographic distance, as was expected (Fig. 6).

### *Species concepts and taxonomic conclusions for* Aspidoscelis

Species concepts present a philosophical quandary (Coyne and Orr, 2004). For example, many modern interpretations [i.e., biological (BSC; Mayr, 1942); evolutionary (ESC; Frost and Hillis, 1990); phylogenetic (PSC; Cracraft, 1983)] offer a mechanistic or philosophical interpretation for the existence of species, yet do not themselves present an explicit operational criterion that can universally define species. The definitions also have fundamental conflicts with regards to the secondary characteristics each relies upon (e.g., reproductive isolation, morphological diagnosability; de Queiroz, 2007).

Given that inferring a direct relationship between genotypic (or ‘haplotypic’) clusters is a questionable approach, we instead relied upon a ‘theory-independent’ definition (the ‘genotypic cluster’ concept: Douglas et al., 2007; Sullivan et al., 2014) wherein clusters are defined as. distinguishable groups of individuals that have few or no intermediates when in contact’ (Mallet, 1995). We used this basis to compare among delimitation hypotheses for *Aspidoscelis tigris*, and how they may (or may not) extend to theoretical species definitions such as the BSC.

An expectation of the ‘…little to no intermediates in contact’ criterion is that individuals from separate genotypic clusters should not be any more genetically similar when sampled in proximity than when sampled from more geographically disparate locations. Such a relationship would suggest continuous gene flow limited only by the ability of individuals to disperse (i.e., ‘isolation-by-distance’ *sensu* Wright, 1940, 1943).

We tested the applicability of this criterion to our species hypotheses using spatial and non-spatial analyses of molecular variance (AMOVA/ SAMOVA), by testing the manner by which each partitioning scheme explained spatial signatures of genetic differentiation (IBD/ Mantel), and with qualitative comparisons of landscape-level genetic differentiation (AIS interpolation). We found the ABGD model (i.e., separation of *A. marmoratus* from all *A. tigris* subspecies) did well with partitioning spatial haplotypic variation, with *A. marmoratus* being highly differentiated (>10% sequence divergence) even from localities sampled <2km from *A. t. punctilinealis* (Fig. 6A). These patterns are consistent with many commonly applied species concepts (e.g. reproductive isolation; per BSC), and given this, the elevated status of *A. marmoratus* is sustained.

We also found limited (albeit, existing) exchange between two major clades within the remaining *A. tigris: A. t. tigris/punctilinealis* versus *A. t. septentrionalis/mundus* (i.e., TAX model). Here, genetic differentiation was again uncorrelated with spatial distances (Fig. 6B), possibly implicating intrinsic or extrinsic factors that may satisfy species-level criteria. Previous estimates place divergence of these groups to be no greater than 5-10 kya, based upon availability of suitable habitat following the Wisconsin glacial period (Zweifel, 1962). If so, then our observed mean corrected genetic distance of 0.058 equates to a mutation rate of 5.8 substitutions per million years, a rate considerably greater than the expected rate of ~0.0045 for ATPase 6, the more rapidly evolving of our two mitochondrial genes (Kumar, 1996; Mueller, 2006).

Although we recognize that numerous factors contribute to mtDNA rate heterogeneity, our rudimentary calculation suggests that subspecies within *A. tigris* may have diverged much earlier than previously thought. Although our results support the continued recognition of *A. t. septentrionalis* as distinct, we deem them insufficient to propose elevation beyond a subspecific level. Instead, we argue for greater sampling at both geographical and genomic scales to properly adjudicate taxonomies within *A. t. tigris*. The latter, while enabling tests of hybridization/ introgression within a more statistically robust framework (e.g., Chafin et al., 2019), would also open avenues for novel species delimitation algorithms to be formulated using unsupervised machine learning approaches (Derkarabetian et al., 2019; Martin et al., 2020; Mussmann et al., 2020).

The CD5 model identified potential cryptic lineages within a clade comprising *A. t. tigris* and *A. t. punctilinealis*, although sample sizes representing this group were limited. Of note, *A. t. tigris* and *A. t. punctilinealis* were not reciprocally monophyletic, however assigning causality of this to hybridization/ intergradation, stochastic lineage sorting, or a lack of phylogenetic signal requires additional data (Chafin et al., 2020). Again, we suggest a more in-depth examination using a larger panel of molecular markers so as to confirm or reject continued recognition of *A. t. tigris* as distinct from *A. t. punctilinealis*, and likewise to adjudicate recognition of *A. t. mundus* from *A. t. septentrionalis*.

### Conclusion

We found that algorithmic species delimitation methods vary substantially in the manner by which they partition hierarchical phylogenetic diversity, particularly when contrasted against the respected overlays of historical biogeography, morphology, ecology, and prior genetic evaluations. By this we mean that some algorithms (e.g., ABGD) yielded groupings which support external evidence of divergence at the ‘species’ level (e.g., according to a BSC-like interpretation), while others (e.g., CD5) yield groupings that could potentially fit various operational species concepts, yet require further evaluation using genomic data. Others (GMYC, PTP) seemingly delimit populations, a conclusion in line with prior evaluations of simulated and empirical datasets. The latter was hypothesized to be impacted by inflated levels of interpopulation divergence, a phenomenon which has been observed in other co-occuring desert fauna subjected to the same historical climatic fluctuation. These observations prompt us to support the elevation of *A. marmoratus*, but additional groups withinthe *A. tigris* complex will requiring additional data to properly adjudicate.

## ACKNOWLEDGMENTS

Samples were collected by BKS and JEC under authority of permits provided by Arizona and New Mexico Game and Fish departments. Collecting methods were approved as part of IACUC protocols for surveying and vouchering lizards (BKS and JEC). We are also indebted to students, post-doctorals, and faculty who have promoted this research: M. R. Bangs, L. Graham, B. A. Levine, B. T. Martin, S. M. Mussmann, Z. D. Zbinden, and especially K.E. Harrell. This research was made possible through generous endowments to the University of Arkansas: A Doctoral Dissertation Fellowship (TKC), the Bruker Professorship in Life Sciences (MRD), and the Twenty-First Century Chair in Global Change Biology (MED), all of which provided salaries and/or research funds for this work. Additional analytical resources were provided by the Arkansas Economic Development Commission (Arkansas Settlement Proceeds Act of 2000) and the Arkansas High Performance Computing Center (AHPCC), and from an NSF-XSEDE Research Allocation (TG-BIO160065) to access the Jetstream cloud. Summer support of field work for JEC was derived from consecutive Endowed Professorships in Science by Opelousas General Hospital of Opelousas, Louisiana.

