## Supplemental Materials for "Taxonomic hypotheses and the biogeography of speciation in the Tiger Whiptail complex (*Aspidoscelis tigris*: Squamata, Teiidae)" (from another version of this preprint: version 1, 2020-10-07)

**Supplementary Materials**

*Supplementary Methods – Species Delimitation*

ABGD (Puillandre et al., 2012) was run using the available web-server graphical interface (bioinfo.mnhn.fr/abi/public/abgd/abgdweb.html) using aligned sequences provided as FASTA. The Kimura 2-parameter (Kimura, 1980) corrected p-distances were used, with the algorithm run with a dist_limit_ range of p_min_=0.001 and p_max_=0.10, and a relative gap width=1.5, and number of bins (N_b_)=40. Analyses were replicated with varying number of recursions of 10, 20, and 40; as well as with larger p_min_ values (0.15, 0.20, 0.25), with no detectable changes in the optimal number of groups delimited.

A second distance-based approach utilized simple clustering using the program CD-HIT, with clustering thresholds of 2% and 5% (corresponding to 98% and 95% identity, respectively). Global sequence identity was computer (-G 1), with a word size of 10 (-n 10), an alignment bandwidth of 20 (-b 20), and with default alignment coverage parameters as implemented using the CD-HIT Suite web server graphical interface (weizhong-lab.ucsd.edu/cdhit-web-server/cgi-bin/index.cgi). Sequences were also clustered to the best clustering meeting the identity threshold (-g 1).

Groups were also delimited within the context of phylogenetic relationships in multiple ways. The first, termed ‘ROS’, required groups to be: have acceptable nodal supports (bootstrap proportion; BP>0.7), and pass the Rosenberg (2007) test of reciprocal monophyly, as implemented using the R package Spider (Brown et al., 2012), and using a p-value threshold of 0.05.

GMYC (Pons et al., 2006) was run using the Splits R package (Fujisawa et al., 2013; R Core Team, 2020), using the an ultrametric tree (see ‘Phylogenetic inference’ in main document) and a single-threshold model and default scaling parameters. PTP (Zhang et al., 2013) was computed using the Bayesian implementation provided by the bPTP web server (species.h-its.org) using 100,000 MCMC iterations, with 10% burn-in and retaining every 100^th^ sample. The analysis was replicated multiple times to assess convergence.

**Supplementary Figures and Tables**

Table S1: Analysis of molecular variance (AMOVA) for clade assignments under 6 *de novo* species delimitation hypotheses and currently accepted taxonomic assignments.

| Level | Model | df | Percentage of Variation |
| --- | --- | --- | --- |
| *Among clades within tigris* | D ~ PHY | 7 | 87% |
|  | D ~ ABGD | 1 | 72% |
|  | D ~ TAX | 2 | 79% |
|  | D ~CD5 | 4 | 80% |
|  | D ~ CD2 | 13 | 90% |
|  | D ~ GMYC | 11 | 89% |
|  | D ~ PTP | 15 | 90% |
| *Among individuals within clades* | D ~ PHY | 195 | 13% |
|  | D ~ ABGD | 201 | 28% |
|  | D ~ TAX | 200 | 21% |
|  | D ~CD5 | 198 | 20% |
|  | D ~ CD2 | 189 | 10% |
|  | D ~ GMYC | 191 | 11% |
|  | D ~ PTP | 187 | 10% |

Table S2: Hierarchical analysis of molecular variance (3-way AMOVA) for species delimitation models, reporting among-clade variance, and among- versus within- subclade genetic variance calculated for subclades respective to the ABGD model.

| Level | Model | df | Percentage of Variation |
| --- | --- | --- | --- |
| *among ABGD clades within tigris* | D ~ ABGD/PHY | 1 | 56% |
|  | D ~ ABGD/TAX | 1 | 50% |
|  | D ~ ABGD/CD5 | 1 | 51% |
|  | D ~ ABGD/CD2 | 1 | 56% |
|  | D ~ ABGD/GMYC | 1 | 56% |
|  | D ~ ABGD/PTP | 1 | 56% |
| *Among subclades within ABGD* | D ~ ABGD/PHY | 6 | 35% |
|  | D ~ ABGD/TAX | 2 | 34% |
|  | D ~ ABGD/CD5 | 3 | 34% |
|  | D ~ ABGD/CD2 | 12 | 37% |
|  | D ~ ABGD/GMYC | 10 | 37% |
|  | D ~ ABGD/PTP | 14 | 37% |
| *Among individuals within subclades* | D ~ ABGD/PHY | 195 | 9% |
|  | D ~ ABGD/TAX | 200 | 16% |
|  | D ~ ABGD/CD5 | 198 | 14% |
|  | D ~ ABGD/CD2 | 189 | 7% |
|  | D ~ ABGD/GMYC | 191 | 7% |
|  | D ~ ABGD/PTP | 187 | 7% |

Figure S1: Spatial distribution of group membership under alternative species delimitation hypotheses, with individuals of the *Aspidoscelis tigris* complex colored according to clade membership (see Fig. 1B).

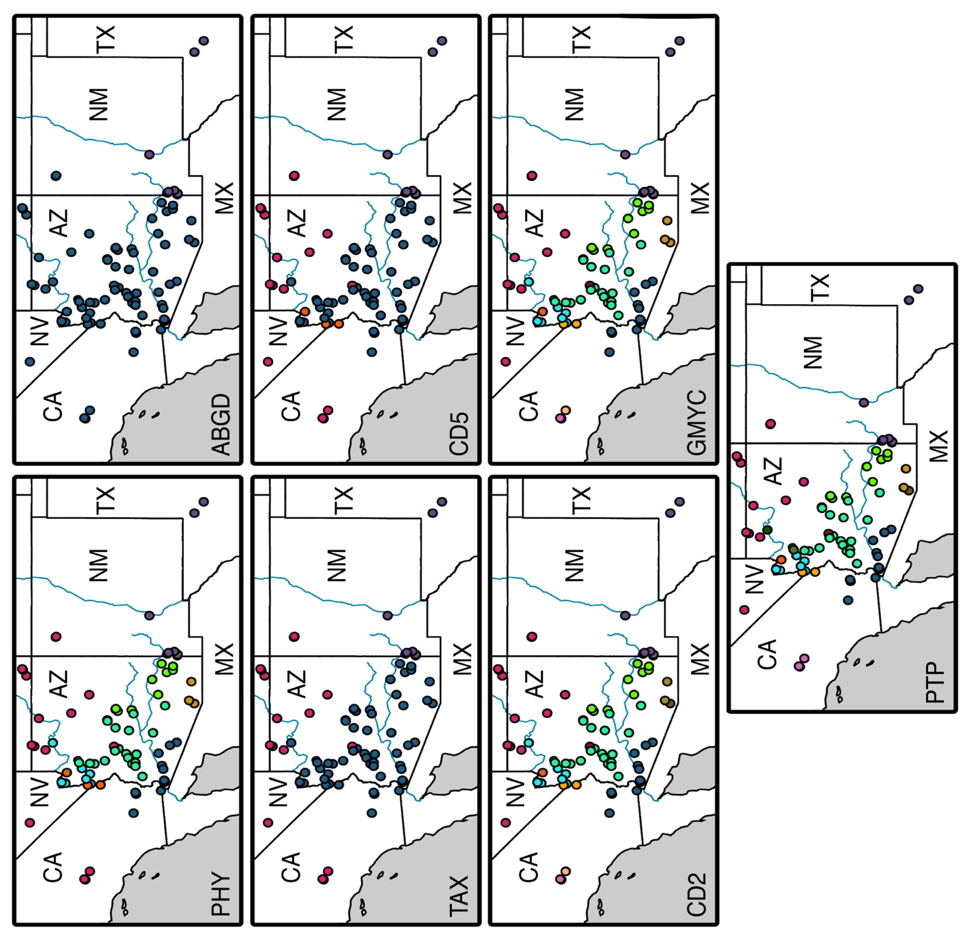
